# Scale-Specific Viscoelastic Characterization of Hydrogels: Integrated AFM and Finite Element Modeling

**DOI:** 10.1101/2025.07.02.662292

**Authors:** N. Fertala, K. Uhlmann, E. Grigoryev, P. Seth, J. Friedrichs, C. Werner, D. Balzani

## Abstract

Viscoelastic hydrogels mimic the dynamic mechanical properties of native extracellular matrices, making them essential for biomedical applications. However, characterizing their scale-dependent mechanical properties remains challenging, despite their critical influence on cell-material interactions and biomaterial performance. Here, we present an integrated experimental-computational approach to quantify and model the viscoelastic behavior of interpenetrating polymer network hydrogels across micro– and macro-scales. Atomic force microscopy-based stress relaxation tests revealed that microgels exhibit rapid, localized relaxation, while macroscopic bulk gels displayed prolonged relaxation dominated by poroelastic effects. Finite element simulations accurately replicated experimental conditions, enabling the extraction of key parameters: instantaneous elastic modulus, relaxation modulus, and relaxation time constant. We further developed a novel analytical model to predict viscoelastic parameters from experimental data with minimal error (< 5%), significantly streamlining characterization. Our findings highlight the necessity of scale-specific mechanical analysis and provide a robust platform for designing biomaterials with tailored viscoelasticity for tissue engineering and regenerative medicine.

## Introduction

Hydrogels have transformed biomedical engineering by mimicking the dynamic, water-rich environment of native extracellular matrices (ECM) through tunable, three-dimensional polymer networks derived from natural or synthetic sources [1–3]. These materials replicate the porous architecture and biochemical signaling of tissues, enabling physiologically relevant models that surpass traditional two-dimensional cultures. As such, hydrogels are integral to advances in targeted drug delivery [4–7], cell encapsulation [8,9], tissue engineering [10–13], biosensing, and mechanobiology [14–18].

A key advantage of hydrogels is their dual tunability in both chemical composition and structural scale. Bulk hydrogels typically serve as robust scaffolds for tissue regeneration, promoting cell adhesion, proliferation, and differentiation, while enabling advanced applications such as 3D bioprinting and organ-on-a-chip platforms [19–24]. In contrast, microscale hydrogel particles, known as microgels (ranging from 0.1 to 100 µm in diameter), excel in cell encapsulation, controlled drug release, and biosensing [25,26], with recent studies demonstrating their ability to guide endothelial morphogenesis through electrostatic microenvironment modulation [27].

Despite their growing utility, microgel mechanics are often extrapolated from bulk gel measurements, assuming that identical formulations yield consistent behavior across scales. However, significant discrepancies exist between micro– and macroscale properties of the same material [28,29]. These differences critically impact biological performance, microgel stiffness and stress relaxation directly affecting cell proliferation, differentiation, and mechanotransduction [30–35]. Inaccurate characterization thus risks compromising biomedical outcomes.

Recent advances in hydrogel technology focus on viscoelastic systems that more accurately recapitulate the time-dependent mechanical behavior of living tissues [36–40]. Unlike purely elastic materials, viscoelastic hydrogels resist immediate deformation and dissipate energy under load, mimicking ECM remodeling during physiological processes such as wound healing [41–43]. However, characterizing these dynamics – particularly at microscales – remains challenging. While bulk rheometry captures macroscale viscoelasticity, probing microgels demands specialized techniques such as atomic force microscopy (AFM)–based nanoindentation [44,45]. Yet, standard stress-relaxation tests under constant strain often oversimplify the complex time-dependent soft material responses, leading to inaccurate estimates of relaxation time and creep compliance.

To address these challenges, we employ previously established viscoelastic interpenetrating polymer network (IPN) hydrogels [46] as model systems to investigate correlations in viscoelastic parameters across multiple scales. These hydrogels feature a dual-network architecture: (1) a covalent network formed through Michael-type addition between thiolated four-armed poly(ethylene glycol) (starPEG) and maleimide-functionalized sulfated glycosaminoglycan heparin, providing structural integrity and elastic behavior; and (2) a physical network arising from reversible electrostatic interactions between heparin and heparin-binding peptides conjugated to starPEG, imparting a time-dependent viscous response (**Figure 1A**) [47,48]. This design allows precise control over both stiffness and stress relaxation behavior, making these IPN hydrogels ideal for exploring scale-dependent viscoelastic phenomena.

**Figure 1:**
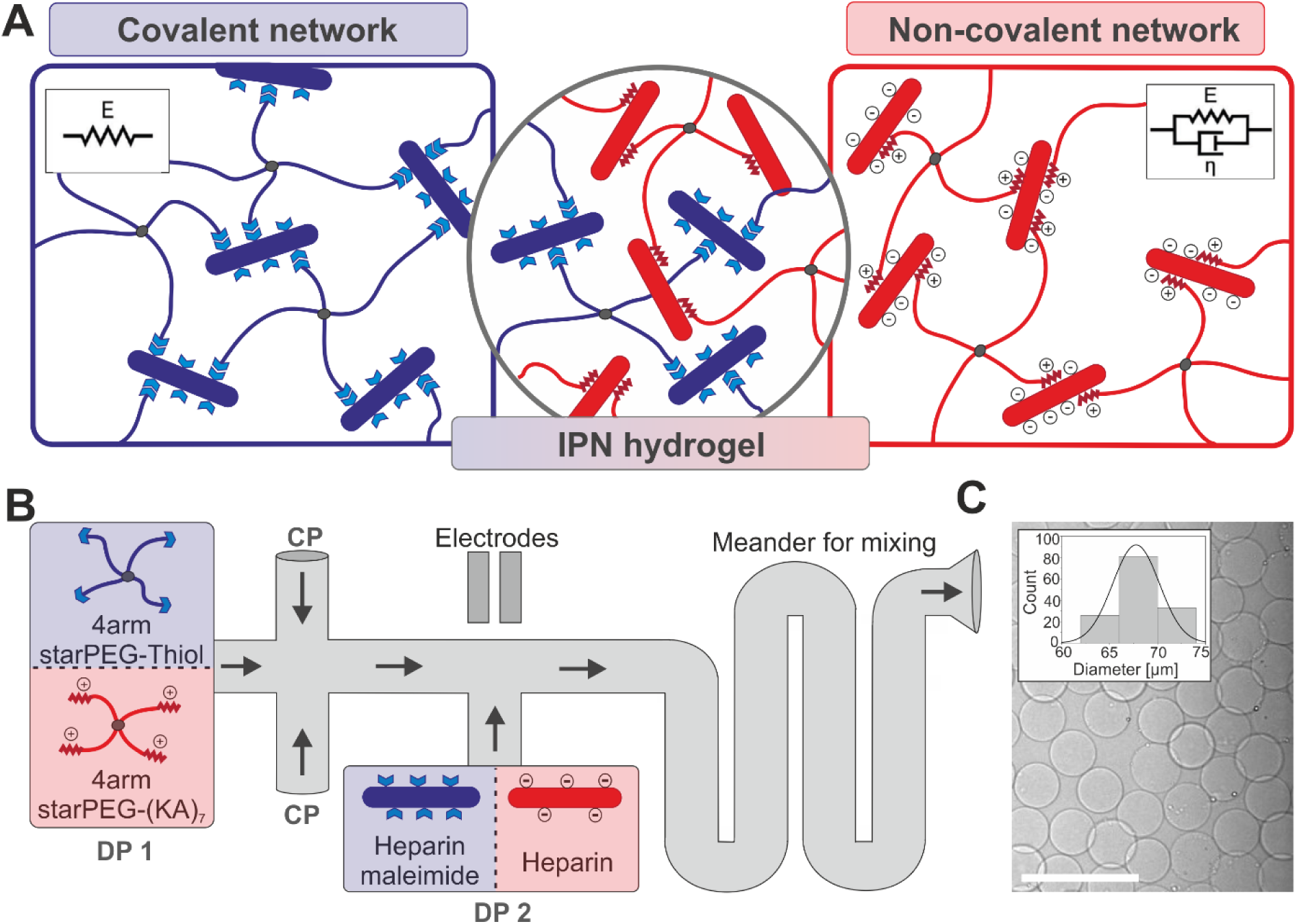
Composition and fabrication of viscoelastic IPN microgels. (**A**) Schematic representation of the viscoelastic interpenetrating polymer network (IPN) hydrogel platform. The hydrogel consists of two networks: a covalent network (blue) formed by a Michael-type addition reaction between thiolated starPEG and maleimide-functionalized heparin provides robust elastic properties. Concurrently, a physically crosslinked network (red) formed via reversible electrostatic interactions between sulfated heparin and starPEG conjugated with the heparin-binding peptide (KA)₇ imparts viscous properties and controlled stress relaxation [46]. **(B)** Microfluidic fabrication of IPN microgels [53]. A microfluidic device equipped with integrated electrodes enables rapid and precise mixing of the continuous phase (CP) of fluorinated oil to which a surfactant is added for droplet stabilization and the precursor solutions. Droplets containing PEG-based precursors (dispersed phase 1 – DP1) and heparin-based precursors (dispersed phase 2 – DP2) are generated separately, merged at a T-junction through electric field-induced droplet fusion, and uniformly mixed within a downstream meandering channel to form monodisperse microgels. Arrows indicate fluid flow directions. **(C)** Brightfield microscopy image of the resulting viscoelastic IPN microgels. The inset shows the size distribution of the microgels (mean = 67.8 ± 2.4 µm, n = 140). Scale bar: 250 µm.

By integrating AFM-based stress relaxation tests with finite element (FE) simulations, we dissected scale dependent viscoelasticity, revealing that bulk gels exhibit prolonged poroelastic relaxation – where water migration through an extensive pore network contributes to a viscous response [49–52], while microgels display rapid, localized relaxation. These scale-dependent differences underscore that inferring microgel properties from bulk gel measurements can lead to substantial discrepancies in key parameters such as relaxation time and creep compliance.

We further developed an analytical model to predict the viscoelastic properties of microgels with error margins below 5%, streamlining characterization. This approach directly correlates AFM measurements with optimized material parameters, offering unprecedented accuracy in capturing both elastic and viscous contributions to hydrogel behavior. Ultimately, our approach bridges micro-macro mechanical discrepancies, offering critical insights for biomaterial design.

## Results and Discussion

### Formation of Viscoelastic Microgels and Bulk Hydrogels

Viscoelastic microgels and bulk hydrogels were fabricated using an established cell-instructive interpenetrating polymer network (IPN) hydrogel platform (**Figure 1A**) [46]. This system combines two distinct networks: (1) a covalent network formed through Michael-type addition between thiolated star-shaped polyethylene glycol (starPEG) and maleimide-functionalized heparin that provides elastic properties; and (2) a physical network of reversible electrostatic interactions between heparin-binding peptides ((KA)₇) conjugated to starPEG and negatively charged heparin sulfate groups, imparting viscous properties and enabling controlled stress relaxation. By adjusting the ratio of these two networks, the viscoelastic profile of the hydrogel can be precisely tuned to mimic natural tissue mechanics.

For the first time, monodisperse microgels were fabricated from this IPN system using microfluidic techniques with integrated electrodes, enabling rapid and controlled mixing of hydrogel precursors via emulsion droplet coalescence (**Figure 1B**) [53]. Droplets of the PEG-based precursor solution (DP 1) were generated at a flow-focusing junction using a fluorinated oil continuous phase (CP) stabilized by surfactant. These droplets were then guided to merge with heparin-based precursor droplets (DP-2) at a T-junction, where an applied electric field (50 kV) triggered rapid coalescence by destabilizing the water-oil interface. Subsequent mixing in a meandering channel facilitated uniform gelation, producing highly monodisperse microgels (diameter = 67.8 ± 2.4 μm, **Figure 1C**).

### Mechanical Characterization and FE Simulations Reveal Scale-Dependent Behavior

To investigate the scale-dependent mechanical properties of the IPN hydrogel, we conducted AFM-based nanoindentation experiments on both microgels and bulk hydrogels prepared from identical precursor compositions. In these experiments, a probe of defined geometry indents the sample while the cantilever deflection measures the applied force. Fitting the resulting force–indentation curves to the Hertz model yielded the Young’s modulus (*E₀*). Our results revealed that microgels are significantly more compliant than bulk gels, and both formats stiffen with increasing indentation velocity, indicating pronounced time-dependent viscoelastic behavior (**Figure 2A**).

**Figure 2:**
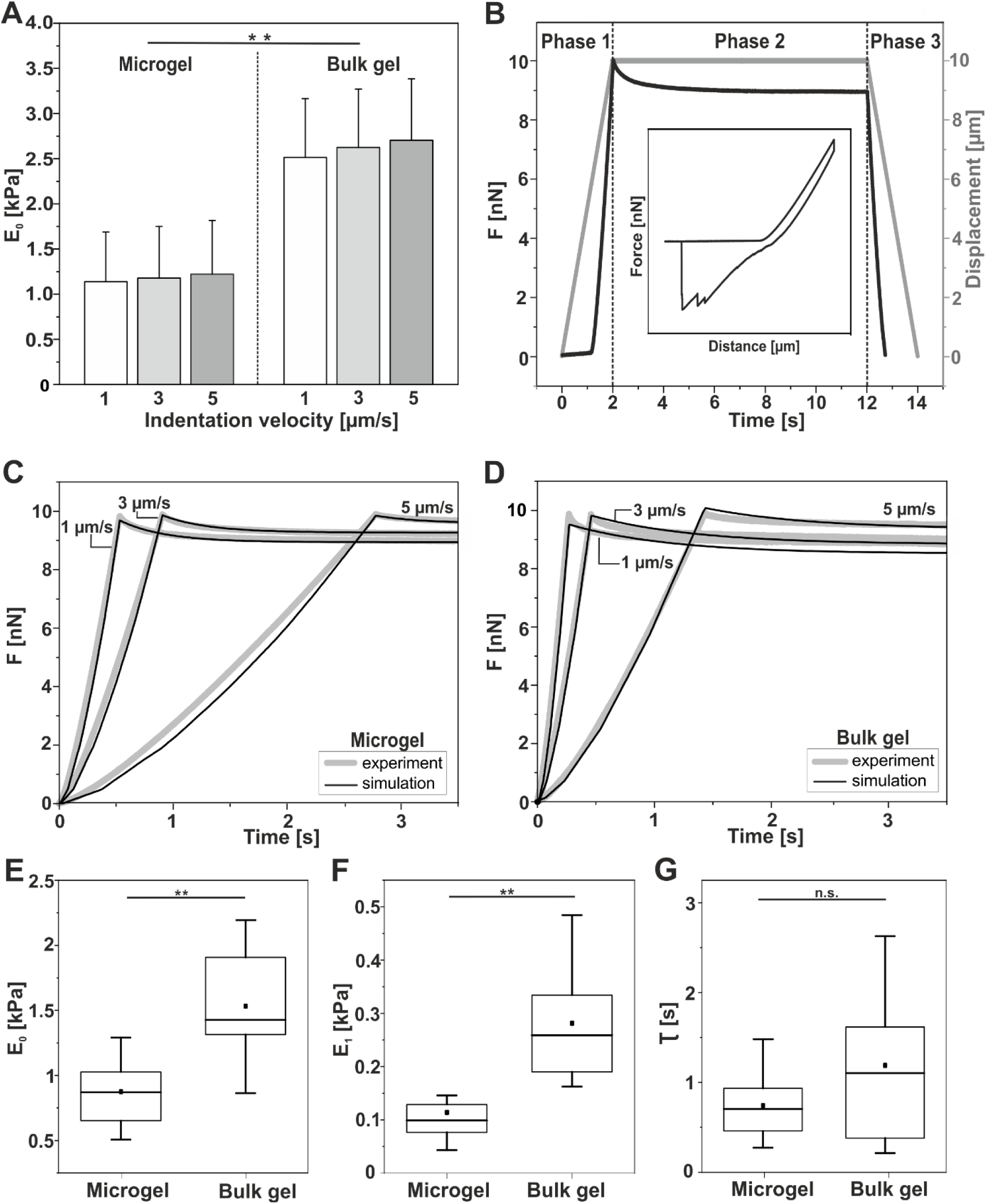
Comparative mechanical characterization of viscoelastic microgels and bulk hydrogels. (**A**) AFM-based nanoindentation analysis comparing elasticity (Young’s modulus, *E₀*) of microgels and bulk gels at varying indentation velocities. Data are presented as mean ± s.d. (n=10). **(B)** Schematic illustration of the three phase AFM-based stress relaxation protocol: Phase 1 (Indentation): The cantilever approaches and indents the sample at constant velocity until a predefined contact force (10 nN) is reached. Phase 2 (Stress Relaxation): The cantilever position is held constant (for 10 s) at maximum displacement, to monitor force decay due to material relaxation. Phase 3 (Retraction): The cantilever is withdrawn from the sample surface. The graph shows representative cantilever height (grey) and contact force (black) profiles over time, with inset illustrating the force–height relationship. **(C, D)** Experimental (grey) and simulated (black) relaxation curves for microgels **(C)** and bulk gels **(D)** across indentation velocities. **(E–G)** FE-derived viscoelastic parameters. Box plots show 25th/75th percentiles (boxes), means (squares), medians (lines), and standard deviation (whiskers). **P < 0.01 (Wilcoxon–Mann–Whitney test).

To further characterize the viscoelastic properties, we performed stress relaxation tests at multiple indentation velocities, following a three-phase protocol (**Figure 2B**). In the first phase, the AFM cantilever approached the sample surface at constant velocity until reaching a predetermined indentation force, inducing instantaneous elastic deformation. Continuous recording of cantilever deflection during this phase provided force–indentation data directly correlated to the material’s instantaneous elastic modulus. These initial measurements serve as the baseline for understanding the load-bearing capabilities of the sample. In the second phase, the cantilever position was held constant at maximum indentation for 10 s, during which the initial stress decays as the polymer network rearranges. This phase yields key viscoelastic parameters – relaxation modulus (*E₁*) and time constant (*τ*) – while the residual stress isolates the purely elastic component (**inset, Figure 2B**). In the third phase, the cantilever was retracted from the sample surface. However, this phase was omitted from quantitative analysis due to adhesion artifacts that obscure intrinsic material behavior.

Both microgels (**Figure 2C**) and bulk gels (**Figure 2D**) demonstrated distinct velocity-dependent relaxation profiles characteristic of viscoelastic materials. To quantitatively extract viscoelastic parameters from these curves, FE simulations replicating the AFM experiments were performed (see Methods). The simulated stress relaxation curves (black lines, **Figure 2C and D**) closely matched experimental data across different indentation velocities, validating the measurement approach and confirming the observed velocity dependence.

However, bulk gels displayed significant deviations during the relaxation phase (Phase 2), particularly at higher indentation velocities, characterized by prolonged and inconsistent relaxation patterns (**Supplementary Figure 1**). This discrepancy likely arises from two competing relaxation mechanisms: (1) intrinsic viscoelasticity arising from polymer network rearrangements, and (2) poroelastic effects caused by fluid migration through the gel’s porous structure during sustained deformation [49,50,54]. The FE simulations, which modeled only the viscoelastic response without accounting for poroelastic effects, consequently showed discrepancies with experimental data during extended relaxation periods.

Bulk gels exhibit prolonged relaxation effects beyond ten seconds, primarily due to poroelastic fluid migration through their extensive pore networks [55,56], as evidenced by non-converging contact forces during indentation. In contrast, microgels exhibited primarily localized viscoelastic relaxation with minimal poroelastic contribution, likely due to their smaller dimensions restricting fluid movement. These differences likely arise from both dimensional effects and fabrication-induced structural variations: bulk gels are formed via active mixing, while microgels rely on diffusion-driven mixing in microfluidic channels, potentially introducing subtle structural variations.

Direct comparison of the FE-derived mechanical parameters revealed significant scale-dependent differences between microgels and bulk hydrogels, despite identical hydrogel composition. Specifically, microgels showed significantly lower Younǵs moduli (*E₀*) compared to bulk gels (**Figure 2E**), along with substantially reduced relaxation moduli (*E_1_*; **Figure 2F**) and shorter relaxation times (*τ*; **Figure 2G**). These findings demonstrate that hydrogel mechanical properties cannot be simply extrapolated across length scales, highlighting the critical importance of scale-specific mechanical characterization for accurate biomaterial design and application.

### Characterization of Material Parameters Using Predefined Equations

While the described optimization procedure accurately determines viscoelastic material parameters, it becomes inefficient when applied to large experimental datasets. To address this limitation, we developed a streamlined analytical approach based on a set of predefined equations that extract key viscoelastic parameters from just four experimental measurements (**Figure 3A**). In addition to these four measurements, the microgel radius (*R*) and the maximum indentation force (*F_1_*, marking the transition between phases 1 and 2) are required. The governing equations are:

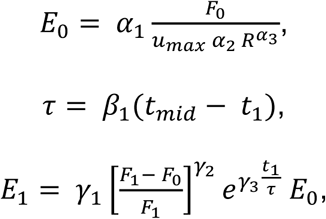

**Figure 3:**
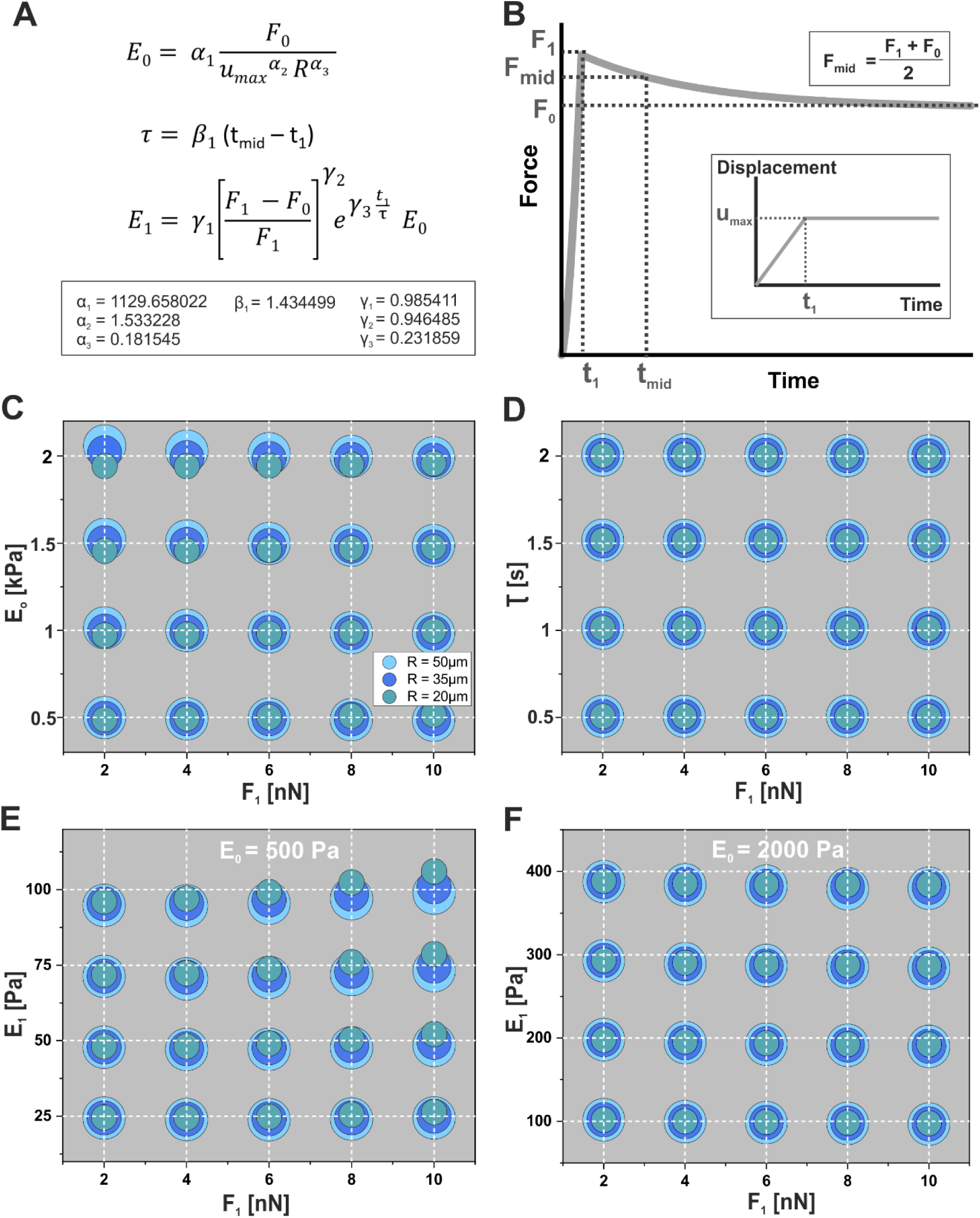
Determination of viscoelastic material parameters and associated errors: (**A**) Predefined equations enable fast estimation of key viscoelastic parameters – fully relaxed Young’s modulus (*E₀*), relaxation modulus (*E₁*), and relaxation time constant (*τ*) – directly from AFM-based stress relaxation data. **(B)** Representative force–time and indentation–time curves illustrating the experimental input parameters required for the calculations shown in (A). **(C–F)** Error analysis comparing estimated parameter values (dots) to actual values (grid intersections) as functions of the maximum contact force (*F₁*) and the microgel radius (*R*): **(C)** Maximum relative errors for Young’s modulus (*E₀*). **(D)** Maximum relative errors for relaxation time (*τ*), observed under conditions of highest relaxation (cantilever velocity *v* = 5 µm/s, *E₀* = 2000 Pa, *E₁* = 400 Pa). **(E, F)** Maximum relative errors for relaxation modulus (*E₁*) at the lower (*E₀* = 500 Pa, **E**) and upper (*E₀* = 2000 Pa, **F**) limits of tested Young’s moduli under conditions of lowest relaxation.

where *F_1_* is the maximum contact force between cantilever and microgel, *F_0_* is the residual contact force at the end of phase 2, *t₁* and *t_mid_* denote the start and midpoint times of phase 2 (with *F_mid_* = (*F_1_* + *F_0_*)/2), and *u_max_* is the maximum indentation depth during phase 2 (**Figure 3B**). The coefficients (α_1_ α_2_, α_3_, β_1_, γ_1_, γ_2_, γ_3_) were optimized across experimentally relevant ranges of microgel radii, maximum contact forces, cantilever velocities, and material parameters (*E_0_, E_1_/E_0_* and *τ*, see Methods).

Using these optimized coefficients, the predefined equations achieved maximum relative errors of 3.1% for *E_0_*, 1.3% for *τ* and 6.0% for *E_1_* across the investigated parameter space. Detailed error analyses (**Figure 3C-F**) illustrate the accuracy of these equations for microgel radii ranging from 20 µm to 50 µm, maximum contact forces from 2 nN to 10 nN, and Young’s moduli up to 2000 Pa. Consistent with experimental observations, the relaxation modulus (*E_1_*) was considered up to 20% of the Young’s modulus (*E_0_*). Specifically, **Figure 3C** shows the relative error in calculated *E₀* (*E₀,cal*) across varying *F₁* and *R*, with the absolute error peaking at *E_0_* = 2000 Pa and *F_1_* = 2 nN. Similarly, the relative error for the relaxation time constant τ remained consistently low (1.3%, **Figure 3D**). The absolute error in calculating E_1_ was most pronounced at Young’s Moduli (*E_0_*) of 500 Pa and 2000 Pa, particularly when the difference between *F_1_* and *F_0_* was minimal, corresponding to the lowest viscosity condition. **Figures 3E and 3F** illustrate these absolute errors in the relaxation modulus (*E_1_*) at *E_0_* = 500 Pa and *E_0_* = 2000 Pa, respectively, at a cantilever velocity *v* = 3 µm/s and *τ* = 0.5 s, varying *F₁* and microgel radii (*R*). Comparable absolute errors for *E₁* were observed at intermediate Young’s moduli (*E₀* = 1000 Pa and 1500 Pa).

Notably, at very low cantilever velocities (i.e. 1 µm/s), the apparent viscosity decreases significantly, causing the predefined equations for *E_1_* to exhibit higher relative errors (up to 30%). Additionally, the equations were specifically optimized for a spherical AFM probe radius of 5 µm. For significantly smaller microgels (< 20 µm) or different cantilever geometries, re-optimization of the coefficients α, β, γ may be necessary. Within the validated parameter ranges, however, these equations provide a rapid, accurate, and straightforward method for extracting viscoelastic material parameters from AFM-based stress relaxation experiments, significantly enhancing evaluation efficiency without significant compromises in accuracy.

## Conclusion

This study introduces an integrated approach for characterizing scale-dependent viscoelastic properties in hydrogels, combining advanced microfluidic fabrication, AFM-based nanoindentation, and computational modeling.

We employed a modular hydrogel system composed of two IPNs, enabling precise and independent control over elasticity and stress relaxation. For the first time, monodisperse microgels were fabricated from this IPN system using microfluidic techniques with integrated electrodes, enabling rapid and controlled mixing of hydrogel precursors via emulsion droplet coalescence. Bulk gels prepared from identical formulations provided a direct basis for comparing viscoelastic properties across different length scales. AFM nanoindentation experiments, supported by FE simulations, revealed that hydrogel mechanics cannot be extrapolated across length scales, highlighting the necessity of scale-specific characterization for accurate biomaterial design.

To facilitate practical implementation, we developed a universal analytical framework that rapidly extracts key viscoelastic parameters – immediate elastic modulus (*E_0_*), relaxation modulus (*E_1_*), and relaxation time (*τ*) – from AFM stress relaxation data. This approach achieves remarkable accuracy (maximum errors: 3.1% for *E₀*, 1.3% for *τ*, 6.0% for *E₁*) across physiologically relevant ranges while eliminating the need for computationally intensive optimization.

This integrated experimental-computational strategy provides a robust platform for designing viscoelastic hydrogels with tailored properties and is particularly valuable for applications in tissue engineering and cellular mechanobiology, where precise control over time-dependent mechanical properties is crucial for guiding cell behavior.

## Materials and Methods

### Hydrogel Components

The IPN hydrogels were prepared from starPEG and heparin precursors. The starPEG components included a commercially sourced starPEG-Thiol (0.83 mM; MW 10,600 g/mol; Polymer Source, Inc.) and an in-house synthesized starPEG-(KA)_7_ (0.28 mM; MW 17,000 g/mol). The heparin components comprised non-functionalized heparin (0.28 mM; MW 14,000 g/mol; Merck KGaA) and an in-house synthesized maleimide-functionalized heparin HM6 (0.83 mM; MW 15,000 g/mol). The viscoelastic properties of the IPN hydrogels were precisely tuned by controlling the network ratio, with the physical (non-covalent) network constituting 25% of the total molar composition and the covalent network comprising the remaining 75%.

### Microfluidic Device Fabrication and Setup Design

Water-in-oil emulsions were generated using custom poly (dimethylsiloxane) (PDMS) microfluidic devices fabricated through standard photolithography and soft lithography techniques [53]. The microfluidic channels were uniformly designed with a width of 50 µm for the dispersed phases, continuous phase, and droplet-transporting channel. The fabrication process began by spin-coating a 3-inch silicon wafer (Siegert Wafer) with SU-8 2025 photoresist (Micro Resist Technology) to create a master mold. Channel patterns were defined using a mask aligner (MJB3, Suess MicroTec) and a printed film mask. After exposure and baking, the uncured photoresist was removed using a developer (mr-Dev 600, Micro Resist Technology). The microchannel structure was then replicated via PDMS replica molding. A mixture of PDMS base and curing agent (Sylgard® 184, Dow Corning) at a 10:1 ratio was degassed using a planetary centrifugal mixer (ARE-250, Thinky), poured over the master mold and cured at 65°C for 2 hours. After demolding, 1.0 mm inlet/outlet ports were punched with a biopsy tool (KAI Medical), and the PDMS channel side was bonded to a glass microscopy slide via oxygen plasma treatment (80Wfor 15 s, MiniFlecto 10, Plasma Technology). To enable electric field-assisted droplet manipulation, electrode microchannels were filled with a low-melting metal alloy (51% In, 32.5% Bi, 16.5% Sn, Indium Corporation of America) to generate localized strong electric fields. Non-electrode microchannels were hydrophobized by injecting a 1% (v/v) solution of (tridecafluoro-1,1,2,2-tetrahydrooctyl)trichlorosilane (Gelest) in Novec 7500 (IoLiTec). During operation, dispersed phases were delivered through 500 µL gastight syringes (Hamilton) and continuous phases via 3 mL disposable syringes (BD Luer-lock), all controlled by high-precision syringe pumps (Pico Plus Elite) connected through PE tubing (0.38 mm ID). The emulsion process was monitored in real-time using an inverted bright-field microscope (Axio Vert.A1, Carl Zeiss) equipped with a high-speed digital camera (Miro C110, Vision Research Inc.).

### Microgel Fabrication

Microgel fabrication was initiated by injecting the first dispersed phase (DP-1), containing 0.83 mM starPEG-Thiol and 0.28 mM starPEG-(KA)_7_, at a flow rate of 50 µL hr^-1^ (**Figure 1B**). Emulsification was achieved using two continuous phases of 2% (w/w) PFPE-PEG-PFPE triblock copolymer surfactant (RAN Biotechnologies) in Novec 7500, delivered at 500 µL hr^-1^. The second dispersed phase (DP-2), containing 0.83 mM heparin-HM6 and 0.28 mM heparin, was introduced through a side channel at matching flow rates, creating paired DP-1/DP-2 droplets. These pairs coalesced into single droplets upon exposure to a 50 kV electric field at integrated electrodes. Subsequent rapid mixing and gelation occurred in a meandering channel, driven by the chemical reaction between the components of DP-1 and DP-2. Microgels were collected every 10 minutes and purified through three washes with a 20% (v/v) 1H,1H,2H,2H-perfluoro-1-octanol (PFO, Sigma-Aldrich) in Novec 7500, before being transferred to PBS buffer.

### Bulk Gel Preparation

Preparation of IPN bulk gels involved two separate precursor solutions: (1) a starPEG solution prepared by combining starPEG-(KA)_7_ with starPEG-Thiol, and (2) a heparin solution prepared by combining non-functionalized heparin with heparin HM6. These solutions were mixed at a 25:75 molar ratio of non-covalent to covalent networks – matching the microgel formulation – and deposited as droplets (20 µL) in Petri dishes filled with PBS. After crosslinking, the resulting hydrogel droplets measured approximately 5 mm in diameter and 1 mm in height.

### Mechanical Characterization of Microgels and Bulk Gels

Mechanical properties were quantified using a NanoWizard 4 AFM (Bruker) coupled with an inverted optical microscope (Observer A1, Zeiss). A tipless cantilever (PNP-TR-TL, NanoWorld; k = 0.08 N/m) fitted with a 10 µm colloidal force probe (Microparticles GmbH) was employed for both indentation and stress relaxation measurements. The cantilever was calibrated before each experiment using the thermal noise method [57] with all measurements conducted in PBS at room temperature.

For indentation testing, the colloidal probe was positioned over the center of individual microgels or bulk gels. Force-distance curves were recorded at three approach/retract velocities (1, 3 and 1 µm/s). A constant contact force of 2 nN was maintained, resulting in typical indentation depths of approximately 1 µm. The Young’s modulus was determined by fitting the approach segment of the force–distance curve to the Hertz model for a spherical indenter. Microgels analysis incorporated a double-contact correction to account for additional deformation from the counter pressure at the microgel’s underside [58–60]. All data processing was performed using the AFM’s native software with custom analysis algorithms.

Stress relaxation measurements followed a three-step protocol (**Figure 2B**). In the indentation phase (Phase 1), the cantilever advanced at controlled velocities (1, 3, or 5 µm/s) until reaching a 10 nN loading force. The subsequent hold phase (Phase 2) maintained constant displacement for 10 seconds while monitoring force decay, allowing characterization of time-dependent viscoelastic behavior. Finally, in the retraction phase (Phase 3), the cantilever was withdrawn from the sample surface.

### Determination of FE Models

To computationally replicate the AFM experiments, FE models were developed representing both microgel and bulk gel geometries. Microgels were modelled as spheres (**Supplementary Figure 2A**), while bulk gels were approximated as cylinders with 1000 µm height to match experimental conditions (**Supplementary Figure 2B**). Both geometries incorporated rotational symmetry to optimize computational efficiency.

A mesh convergence study established the optimal element density for accurate parameter determination (**Supplementary Figure 2C**). The microgel model comprised 23,690 elements (24,130 degrees of freedom), while the bulk gel model used 14,714 elements (15,080 degrees of freedom). Triangular elements with quadratic shape functions were employed throughout, with localized mesh refinement in the bulk gel’s contact region to resolve steep stress gradients. The AFM indenter was modeled as a rigid 10 µm sphere with frictionless contact assumptions.

### Finite Element Simulation-Based Analysis of Viscoelastic Material Properties

FE simulations were employed to analyze the viscoelastic behavior of microgels and bulk gels, coupled with an optimization procedure to determine material parameters that best fit experimental data. Given the significant displacements observed during AFM indentation, the material model was based on finite strain theory. The viscoelastic material behavior was described using a combination of a Neo-Hookean model and a single-term Prony series. The Neo-Hookean model captured the nonlinear elastic response at large deformations, while the Prony series component accounted for the time-dependent viscous response. This approach is equivalent to a generalized Maxwell model with a single spring-dashpot element, commonly used in relaxation experiments [61–63].

The time-dependent Young’s modulus *E(t)* is expressed in one dimension as:

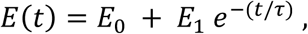

where *E*_0_is the fully relaxed Young’s modulus, *E*_1_represents the initial viscous contribution, and τ is the relaxation time constant. The instantaneous (unrelaxed) modulus is given by *E*_*u*_ = *E*_0_ + *E*_1_, and τ can be interpreted as τ = η/*E*_1_, with η being the dashpot viscosity. To account for the nearly incompressible nature of the hydrogels, the Poisson’s ratio was set to *ν* = 0.45 [14,59].

Parameter optimization proceeded in two sequential phases:

1. Relaxed Modulus (*E_0_*): The fully relaxed behavior was determined using phase 2 of the experiment, during which the indenter was held at the maximum displacement (*u*_*max*_) for 10 seconds (**Figure 2B**).
2. Viscous Parameters (*E_1_* and τ): These parameters were optimized using the stress relaxation data from phase 2, where the displacement remained constant, and the stress decay was governed solely by viscous behavior.

This two-step approach minimized inaccuracies caused by potential nonlinear elastic effects not fully captured by the Neo-Hookean model during the initial loading phase.

### FE Simulation and Parameter Optimization

FE simulations were performed in Abaqus (Dassault Systèmes. Abaqus 2024: Unified FEA Software), a commercial finite element software, with optimization implemented in Python (Python Software Foundation (2023)) using the “mystic” library, which combines evolutionary strategies with gradient-based methods.

#### Elastic Modulus (*E₀*) Optimization

*E₀* was optimized using the maximum indenter displacement (*u*_*max*_) from the experiments as the boundary condition. Since experiments were conducted at three indenter velocities (*v₁* = 1 µm/s, *v₂* = 3 µm/s, and *v₃* = 5 µm/s), three separate simulations were executed for each candidate *E_0_* value. In these simulations, the material was modeled using a Neo-Hookean model (without viscosity). The simulated contact force (*F_sim_)* at the end of the relaxation phase (phase 2) was compared with the corresponding experimental force (*F_exp_)*. The first objective function *Z*_1_ of the optimization was defined as:

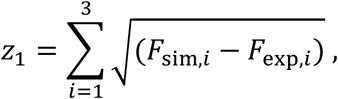

where *F*_sim,*i*_ and *F*_exp,*i*_ are the contact forces at the end of phase 2 for the three velocities (*v*_1_, *v*_2_ and *v*_3_).

#### Optimizing the Viscous Parameters (*E₁* and *τ*)

The viscous parameters (*E₁* and *τ*) were optimized by capturing the full force–time response during phases 1 and 2. In these simulations, the experimental displacement– time profile was imposed as the boundary condition. The simulated force–time curves (*F_sim_)* were compared with the experimental curves (*F_exp_)* at all time steps across the three velocities. The second objective function (z_2_) was defined as:

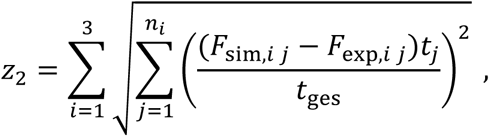

where *i* corresponds to the different velocities (*v*_*i*_) and *n*_*i*_ is the total number of time steps of the simulation in phase 2. Consequently, *F*_sim,*i* *j*_ are the contact forces at the corresponding time step at the time *t*_*j*_ and *F*_exp,*i* *j*_ is the contact force in the experiments, which was linearly interpolated from the experimental data for the time *t*_*j*_. The factor (*t*_*j*_⁄*t*_ges_) with *t*_ges_ = 10s was used as weighting factor for a more reliable optimization result.

### Optimization of Material Parameter Equations

A mathematical model was constructed to estimate *E_0_, E_1_* and *τ* (**Figure 3A**) and the coefficients in these material parameter equations were optimized over the data range specified in **Supplementary Figures 3-6**. To achieve this, 1,920 simulations replicating the AFM mechanical experiments were performed to fine-tune the coefficients α_1_ α_2_, α_3_, β_1_, γ_1_, γ_2_, γ_3_. Each simulation tested a unique combination of the following variables:

- Elastic modulus (*E_0_*): 500, 1000, 1500, 2000 Pa
- Microgel radius (*R*): 20, 35, 50 µm
- Maximum contact force (*F_1_*): 2, 4, 6, 8, 10 nN
- Ratio *E_1_/E_0_*: 0.05, 0.1, 0.15, 0.2
- Relaxation time constant (*τ*): 0.5, 0.1, 0.15, 0.2 s
- Indenter velocity (*v*): 3 µm/s, 5 µm/s.

The radius of the indenter cantilever was fixed at 5 µm.

The prediction of the purely elastic modulus (*E_0_*) was based on the relaxed contact force (*F_0_*) and the microgel radius (*R*). Since only viscoelastic materials were considered, *F_0_* was not predefined but varied with *F_1_, E_1_, τ* and *v*. Error plots were generated for the conditions *E_1_/E_0_* = 0.05, *τ* = 0.5 s, and *v* = 3 µm/s, representing the lowest viscosity and largest absolute errors. For each simulation, the values of *F_0_, u_max_, R, F_1_, t_1_* and *t_mid_* were extracted and used to predict *E_0_, τ* and *E_1_.* The relative error for each simulation was calculated as:

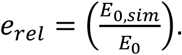

The optimization process adjusted the parameters to minimize the maximum relative error across all simulations.

## Acknowledgements

We gratefully acknowledge Julian Thiele (Leibniz Institute of Polymer Research Dresden, Germany) for insightful discussions and valuable input. This work was supported by the Deutsche Forschungsgemeinschaft (DFG, German Research Foundation) under project number BA 2823/19-1.

## Conflict of Interest

The authors declare no conflict of interest.

## Supplementary Information

**Supplementary Figure 1:**
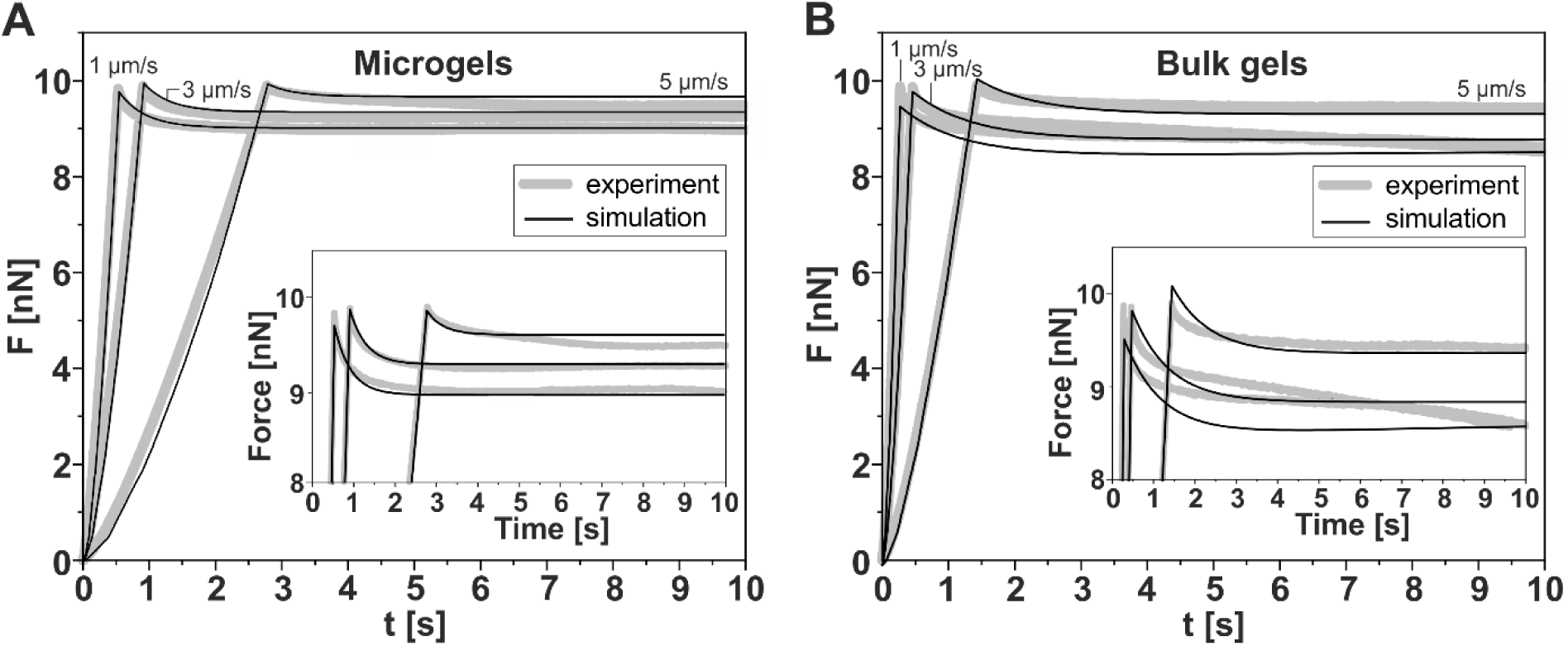
Complete stress relaxation phase of microgels and bulk gels. **(A)** Experimental force–time curves (grey lines) and corresponding FE simulation curves (black lines) for microgels at various indentation velocities, showing the full 10-second holding period of phase 2. **(B)** Equivalent data for bulk gels, where the force–time curves reveal inconsistent and prolonged relaxation behavior, attributable to poroelastic effects during the holding phase. Insets provide enlarged views of phase 2 for both microgels and bulk gels, highlighting the detailed relaxation dynamics at the probe–sample interface.

**Supplementary Figure 2:**
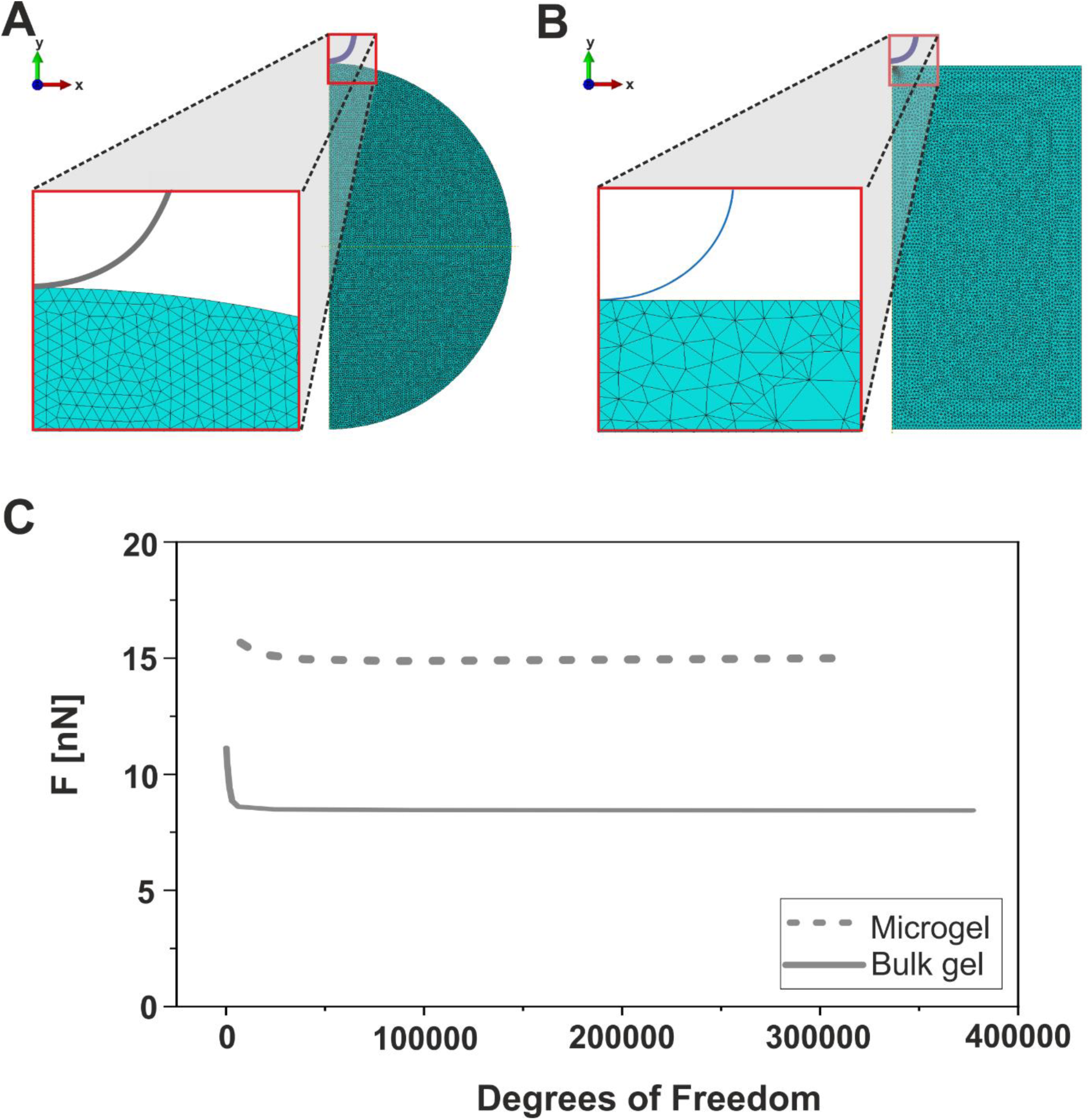
FE Model Geometry and Mesh Convergence Analysis. FE model geometry and mesh used for nanoindentation simulations of microgels **(A)** and bulk gels **(B)**. Insets (red boxes) in both panels provide enlarged views of the contact region between the spherical AFM probe and the sample surface, highlighting local mesh refinement. **(C)** Mesh convergence analysis showing the relationship between computed contact force and increasing degrees of freedom for microgels (dashed line) and bulk gels (solid line), demonstrating that further mesh refinement yields negligible changes in the simulated contact force, thus confirming convergence.

**Supplementary Figure 3:**
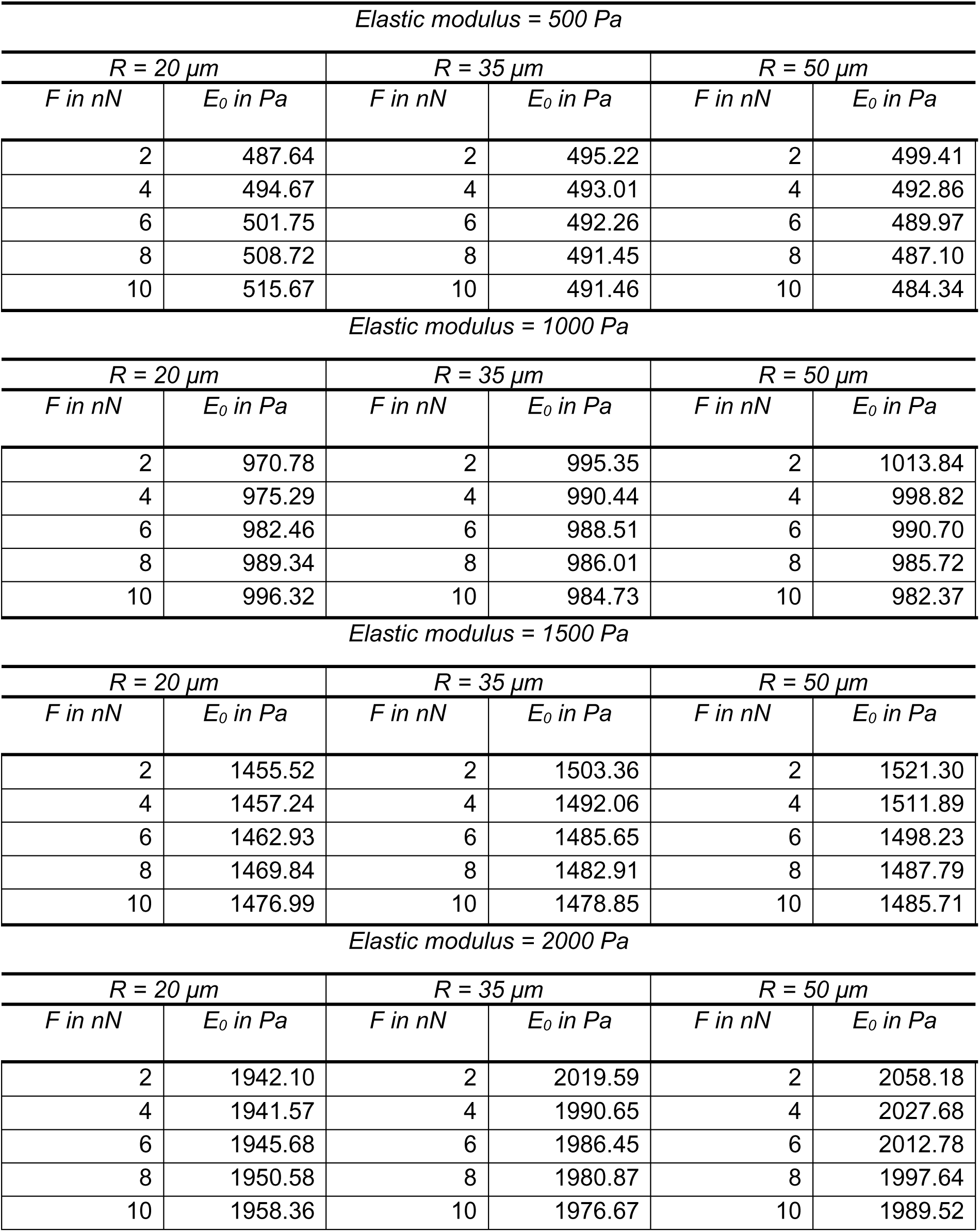
Estimated values of the Young’s modulus (*E₀*) compared to actual values, analyzed as a function of the predefined maximum contact force (*F*) and microgel radius (*R*).

**Supplementary Figure 4:**
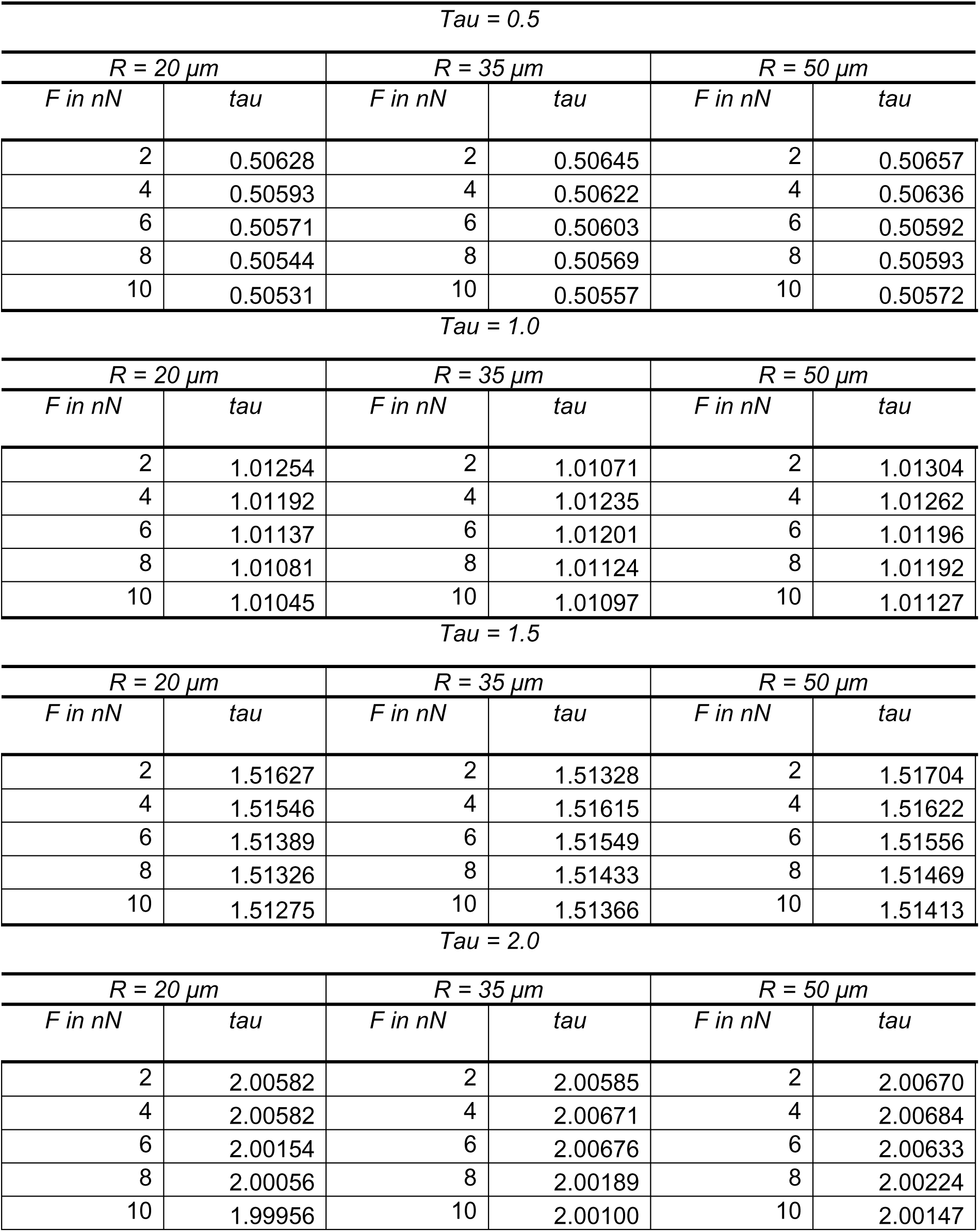
Estimated values of the relaxation time (*τ*) compared to actual values, analyzed as a function of the predefined maximum contact force (*F*) and the radius (*R*) of the microgel.

**Supplementary Figure 5:**
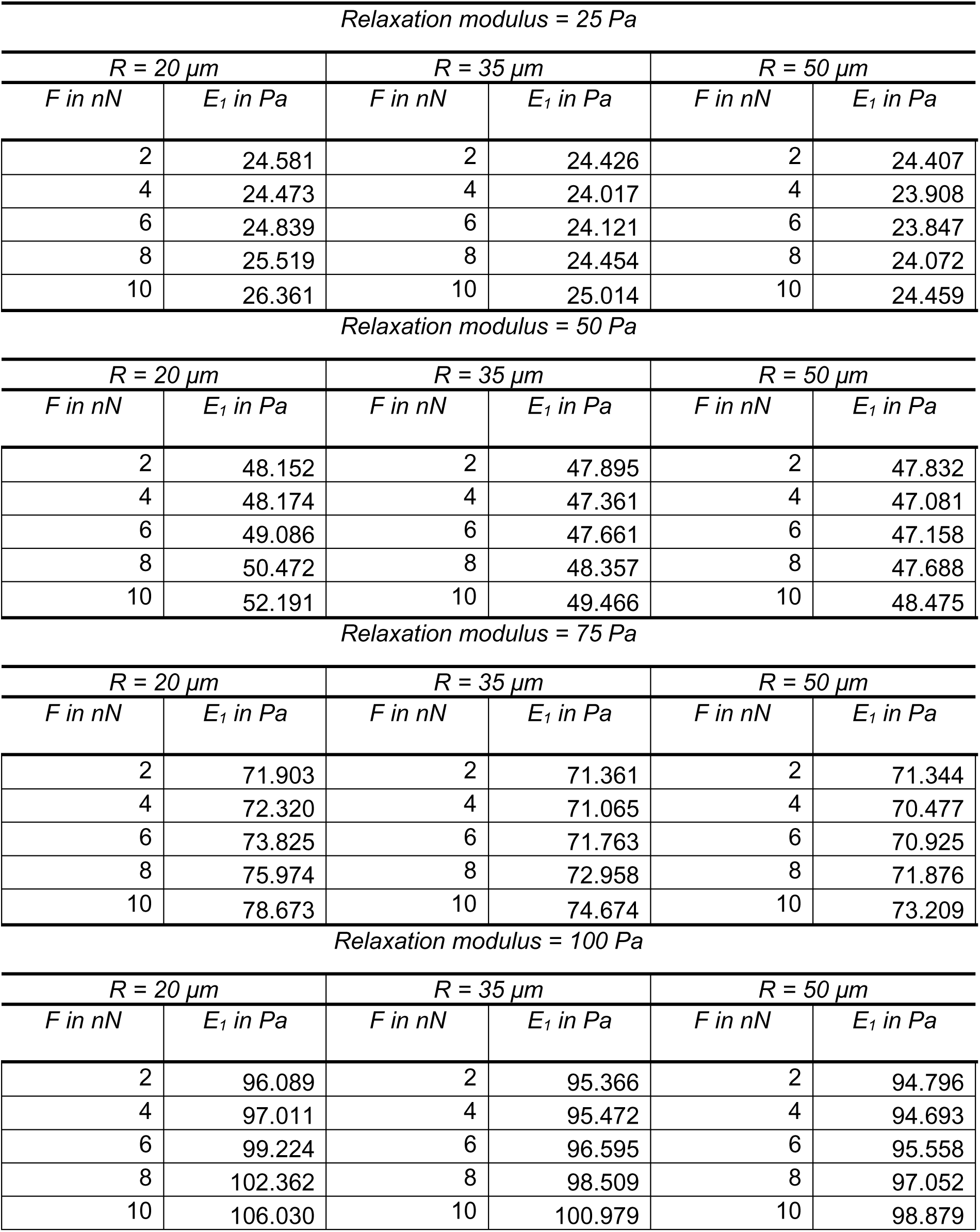
Estimated values for the relaxation modulus (*E₁*) at the lower boundary (with *E₀* = 500 Pa) compared to actual values, analyzed as a function of the predefined maximum contact force (*F*) and the radius (*R*) of the microgel.

**Supplementary Figure 6:**
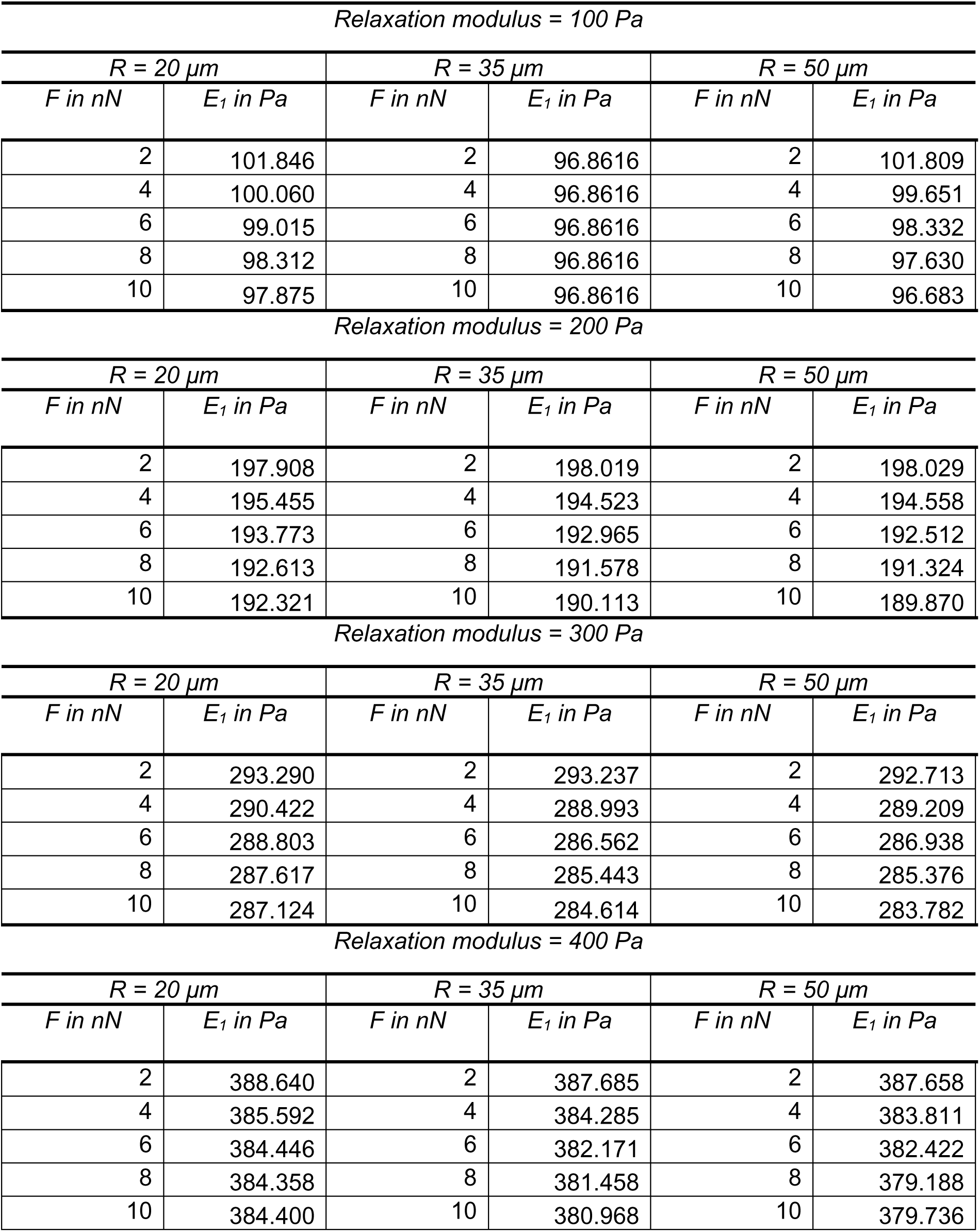
Estimated values for the relaxation modulus (*E₁*) at the upper boundary (with *E₀* = 2000 Pa) compared to actual values, analyzed as a function of the predefined maximum contact force (*F*) and the radius (*R*) of the microgel.

## Notes

### Competing Interest Statement

The authors have declared no competing interest.

## References

[1] Drury JL, Mooney DJ. Hydrogels for tissue engineering: Scaffold design variables and applications. Biomaterials 2003;24:4337–51. 10.1016/S0142-9612(03)00340-5.

[2] Tibbitt MW, Anseth KS. Hydrogels as extracellular matrix mimics for 3D cell culture. Biotechnol Bioeng 2009;103:655–63. 10.1002/bit.22361.

[3] Hoffman AS. Hydrogels for biomedical applications. Adv Drug Deliv Rev 2012;64:18–23. 10.1016/j.addr.2012.09.010.

[4] Li J, Mooney DJ. Designing *_hydrogels_* for controlled drug delivery. Nat Rev Mater 2016;1:16071. 10.1038/natrevmats.2016.71.

[5] Peppas NA, Hilt JZ, Khademhosseini A, Langer R. Hydrogels in biology and medicine: From molecular principles to bionanotechnology. Adv Mater 2006;18:1345–60. 10.1002/adma.200501612.

[6] Koetting MC, Peters JT, Steichen SD, Peppas NA. Stimulus-responsive hydrogels: Theory, modern advances, and applications. Mater Sci Eng R Reports 2015;93:1–49. 10.1016/j.mser.2015.04.001.

[7] Kühn S, Sievers J, Stoppa A, Träber N, Zimmermann R, Welzel PB, et al. Cell-Instructive Multiphasic Gel-in-Gel Materials. Adv Funct Mater 2020;30. 10.1002/adfm.201908857.

[8] Mohajeri M, Eskandari M, Ghazali ZS, Ghazali HS. Cell encapsulation in alginate-based microgels using droplet microfluidics; A review on gelation methods and applications. Biomed Phys Eng Express 2022;8. 10.1088/2057-1976/ac4e2d.

[9] Xuan L, Hou Y, Liang L, Wu J, Fan K, Lian L, et al. Microgels for Cell Delivery in Tissue Engineering and Regenerative Medicine. Nano-Micro Lett 2024;16:1–42. 10.1007/s40820-024-01421-5.

[10] Wei W, Ma Y, Yao X, Zhou W, Wang X, Li C, et al. Advanced hydrogels for the repair of cartilage defects and regeneration. Bioact Mater 2021;6:998–1011. 10.1016/j.bioactmat.2020.09.030.

[11] Zhang L, Su L, Wu L, Zhou W, Xie J, Fan Y, et al. Versatile hydrogels prepared by microfluidics technology for bone tissue engineering applications. J Mater Chem B 2025:2611–39. 10.1039/d4tb02314e.

[12] Choi H, Choi WS, Jeong JO. A Review of Advanced Hydrogel Applications for Tissue Engineering and Drug Delivery Systems as Biomaterials. Gels 2024;10:1–23. 10.3390/gels10110693.

[13] Patel DK, Jung E, Priya S, Won SY, Han SS. Recent advances in biopolymer-based hydrogels and their potential biomedical applications. Carbohydr Polym 2024;323:121408. 10.1016/j.carbpol.2023.121408.

[14] Träber N, Uhlmann K, Girardo S, Kesavan G, Wagner K, Friedrichs J, et al. Polyacrylamide Bead Sensors for in vivo Quantification of Cell-Scale Stress in Zebrafish Development. Sci Rep 2019;9:1–14. 10.1038/s41598-019-53425-6.

[15] Serwane F, Mongera A, Rowghanian P, Kealhofer DA, Lucio AA, Hockenbery ZM, et al. In vivo quantification of spatially varying mechanical properties in developing tissues. Nat Methods 2016;14:181–6. 10.1038/nmeth.4101.

[16] Campàs O, Mammoto T, Hasso S, Sperling R a, O’Connell D, Bischof AG, et al. Quantifying cell-generated mechanical forces within living embryonic tissues. Nat Methods 2013;11:183–9. 10.1038/nmeth.2761.

[17] Vian A, Pochitaloff M, Yen ST, Kim S, Pollock J, Liu Y, et al. In situ quantification of osmotic pressure within living embryonic tissues. Nat Commun 2023;14:1–10. 10.1038/s41467-023-42024-9.

[18] Parvin N, Kumar V, Joo SW, Mandal TK. Cutting-Edge Hydrogel Technologies in Tissue Engineering and Biosensing: An Updated Review. Materials (Basel) 2024;17. 10.3390/ma17194792.

[19] Zhang XN, Zheng Q, Wu ZL. Recent advances in 3D printing of tough hydrogels: A review. Compos Part B Eng 2022;238:109895. 10.1016/j.compositesb.2022.109895.

[20] Liu H, Wang Y, Cui K, Guo Y, Zhang X, Qin J. Advances in Hydrogels in Organoids and Organs-on-a-Chip. Adv Mater 2019;31:1–28. 10.1002/adma.201902042.

[21] Carvalho V, Gonçalves I, Lage T, Rodrigues RO, Minas G, Teixeira SFCF, et al. 3d printing techniques and their applications to organ-on-a-chip platforms: A systematic review. Sensors 2021;21. 10.3390/s21093304.

[22] Limasale YDP, Fusenig M, Samulowitz M, Atallah P, Sievers J, Dennison N, et al. Glycosaminoglycan Concentration and Sulfation Patterns of Biohybrid Polymer Matrices Direct Microvascular Network Formation and Stability. Adv Funct Mater 2024;2411475:1–14. 10.1002/adfm.202411475.

[23] Sievers J, Mahajan V, Welzel PB, Werner C, Taubenberger A. Precision Hydrogels for the Study of Cancer Cell Mechanobiology. Adv Healthc Mater 2023;12. 10.1002/adhm.202202514.

[24] Fang W, Yang M, Wang L, Li W, Liu M, Jin Y, et al. Hydrogels for 3D bioprinting in tissue engineering and regenerative medicine: Current progress and challenges. Int J Bioprinting 2023;9:207–38. 10.18063/IJB.759.

[25] Xu Y, Zhu H, Denduluri A, Ou Y, Erkamp NA, Qi R, et al. Recent Advances in Microgels: From Biomolecules to Functionality. Small 2022;18. 10.1002/smll.202200180.

[26] Newsom JP, Payne KA, Krebs MD. Microgels: Modular, tunable constructs for tissue regeneration. Acta Biomater 2019;88:32–41. 10.1016/j.actbio.2019.02.011.

[27] Kühn S, Magno V, Zimmermann R, Limasale YDP, Atallah P, Stoppa A, et al. Microgels With Electrostatically Controlled Molecular Affinity to Direct Morphogenesis. Adv Mater 2024;2409731:1–13. 10.1002/adma.202409731.

[28] Roberge CL, Kingsley DM, Cornely LR, Spain CJ, Fortin AG, Corr DT. Viscoelastic Properties of Bioprinted Alginate Microbeads Compared to Their Bulk Hydrogel Analogs. J Biomech Eng 2023;145:1–11. 10.1115/1.4055757.

[29] Rubiano A, Galitz C, Simmons CS. Mechanical Characterization by Mesoscale Indentation: Advantages and Pitfalls for Tissue and Scaffolds. Tissue Eng – Part C Methods 2019;25:619–29. 10.1089/ten.tec.2018.0372.

[30] Schulte MF, Izak-Nau E, Braun S, Pich A, Richtering W, Göstl R. Microgels react to force: mechanical properties, syntheses, and force-activated functions. Chem Soc Rev 2022:2939–56. 10.1039/d2cs00011c.

[31] Morley CD, Ding EA, Carvalho EM, Kumar S. A Balance between Inter– and Intra-Microgel Mechanics Governs Stem Cell Viability in Injectable Dynamic Granular Hydrogels. Adv Mater 2023;35:1–12. 10.1002/adma.202304212.

[32] Li B, Zhang L, Yin Y, Chen A, Seo BR, Lou J, et al. Stiff hydrogel encapsulation retains mesenchymal stem cell stemness for regenerative medicine. Matter 2024:3447–68. 10.1016/j.matt.2024.05.041.

[33] Bauer A, Gu L, Kwee B, Li WA, Dellacherie M, Celiz AD, et al. Hydrogel substrate stress-relaxation regulates the spreading and proliferation of mouse myoblasts. Acta Biomater 2017;62:82–90. 10.1016/j.actbio.2017.08.041.

[34] Cantini M, Donnelly H, Dalby MJ, Salmeron-Sanchez M. The Plot Thickens: The Emerging Role of Matrix Viscosity in Cell Mechanotransduction. Adv Healthc Mater 2020;9. 10.1002/adhm.201901259.

[35] Chaudhuri O, Gu L, Klumpers D, Darnell M, Bencherif SA, Weaver JC, et al. Hydrogels with tunable stress relaxation regulate stem cell fate and activity. Nat Mater 2016;15:326–34. 10.1038/nmat4489.

[36] Chaudhuri O, Gu L, Klumpers D, Darnell M, Bencherif SA, Weaver JC, et al. Hydrogels with tunable stress relaxation regulate stem cell fate and activity. Nat Mater 2016;15:326–34. 10.1038/nmat4489.

[37] Elosegui-Artola A, Gupta A, Najibi AJ, Seo BR, Garry R, Tringides CM, et al. Matrix viscoelasticity controls spatiotemporal tissue organization. Nat Mater 2023;22:117–27. 10.1038/s41563-022-01400-4.

[38] Roth JG, Huang MS, Navarro RS, Akram JT, LeSavage BL, Heilshorn SC. Tunable hydrogel viscoelasticity modulates human neural maturation. Sci Adv 2023;9:1–18. 10.1126/SCIADV.ADH8313.

[39] Wu DT, Jeffreys N, Diba M, Mooney DJ. Viscoelastic Biomaterials for Tissue Regeneration. Tissue Eng – Part C Methods 2022;28:289–300. 10.1089/ten.tec.2022.0040.

[40] Crandell P, Stowers R. Spatial and Temporal Control of 3D Hydrogel Viscoelasticity through Phototuning. ACS Biomater Sci Eng 2023;9:6860–9. 10.1021/acsbiomaterials.3c01099.

[41] Hu Y, Jia Y, Wang S, Ma Y, Huang G, Ding T, et al. An ECM-Mimicking, Injectable, Viscoelastic Hydrogel for Treatment of Brain Lesions. Adv Healthc Mater 2023;12:1–11. 10.1002/adhm.202201594.

[42] Li L, Wang S, Chen Y, Dong S, Zhang C, Liao L, et al. Hydrogels mimicking the viscoelasticity of extracellular matrix for regenerative medicine: Design, application, and molecular mechanism. Chem Eng J 2024;498. 10.1016/j.cej.2024.155206.

[43] Patiño Vargas MI, Martinez-Garcia FD, Offens F, Becerra NY, Restrepo LM, van der Mei HC, et al. Viscoelastic properties of plasma-agarose hydrogels dictate favorable fibroblast responses for skin tissue engineering applications. Biomater Adv 2022;139:212967. 10.1016/j.bioadv.2022.212967.

[44] Cuenot S, Fillaudeau A, Briolay T, Fresquet J, Blanquart C, Ishow E, et al. Poroelastic and viscoelastic properties of soft materials determined from AFM force relaxation and force-distance curves. J Mech Behav Biomed Mater 2025;163. 10.1016/j.jmbbm.2024.106865.

[45] Oevreeide IH, Szydlak R, Luty M, Ahmed H, Prot V, Skallerud BH, et al. On the determination of mechanical properties of aqueous microgels-towards high-throughput characterization. Gels 2021;7:1–18. 10.3390/gels7020064.

[46] Seth P, Friedrichs J, Limasale YDP, Fertala N, Freudenberg U, Zhang Y, et al. Interpenetrating Polymer Network Hydrogels with Tunable Viscoelasticity and Proteolytic Cleavability to Direct Stem Cells In Vitro. Adv Healthc Mater 2024;2402656:1–13. 10.1002/adhm.202402656.

[47] Wieduwild R, Tsurkan M, Chwalek K, Murawala P, Nowak M, Freudenberg U, et al. Minimal peptide motif for non-covalent peptide-heparin hydrogels. J Am Chem Soc 2013;135:2919–22. 10.1021/ja312022u.

[48] Wieduwild R, Krishnan S, Chwalek K, Boden A, Nowak M, Drechsel D, et al. Noncovalent hydrogel beads as microcarriers for cell culture. Angew Chem Int Ed Engl 2015;54:3962–6. 10.1002/anie.201411400.

[49] Moeendarbary E, Valon L, Fritzsche M, Harris AR, Moulding D a, Thrasher AJ, et al. The cytoplasm of living cells behaves as a poroelastic material. Nat Mater 2013;12:253–61. 10.1038/nmat3517.

[50] Berry JD, Biviano M, Dagastine RR. Poroelastic properties of hydrogel microparticles. Soft Matter 2020;16:5314–24. 10.1039/d0sm00191k.

[51] Esteki MH, Alemrajabi AA, Hall CM, Sheridan GK, Azadi M, Moeendarbary E. A new framework for characterization of poroelastic materials using indentation. Acta Biomater 2020;102:138–48. 10.1016/j.actbio.2019.11.010.

[52] Kalcioglu ZI, Mahmoodian R, Hu Y, Suo Z, Van Vliet KJ. From macro-to microscale poroelastic characterization of polymeric hydrogels via indentation. Soft Matter 2012;8:3393. 10.1039/c2sm06825g.

[53] Hauck N, Neuendorf TA, Männel MJ, Vogel L, Liu P, Stündel E, et al. Processing of fast-gelling hydrogel precursors in microfluidics by electrocoalescence of reactive species. Soft Matter 2021;17:10312–21. 10.1039/d1sm01176f.

[54] Caccavo D, Cascone S, Lamberti G, Barba AA. Hydrogels: Experimental characterization and mathematical modelling of their mechanical and diffusive behaviour. Chem Soc Rev 2018;47:2357–73. 10.1039/c7cs00638a.

[55] Islam MR, Oyen ML. A poroelastic master curve for time-dependent and multiscale mechanics of hydrogels. J Mater Res 2021;36:2582–90. 10.1557/s43578-020-00090-5.

[56] Li H, Lian X, Guan D. Crossover behavior in stress relaxations of poroelastic and viscoelastic dominant hydrogels. Soft Matter 2023;19:5443–51. 10.1039/d3sm00592e.

[57] Hutter JL, Bechhoefer J. Calibration of atomic-force microscope tips. Rev Sci Instrum 1993;64:1868–73. 10.1063/1.1143970.

[58] Glaubitz M, Medvedev N, Pussak D, Hartmann L, Schmidt S, Helm CA, et al. A novel contact model for AFM indentation experiments on soft spherical cell-like particles. Soft Matter 2014;10:6732–41. 10.1039/C4SM00788C.

[59] Girardo S, Träber N, Wagner K, Cojoc G, Herold C, Goswami R, et al. Standardized microgel beads as elastic cell mechanical probes. J Mater Chem B 2018;6:6245–61. 10.1039/C8TB01421C.

[60] Dokukin ME, Guz N V., Sokolov I. Quantitative study of the elastic modulus of loosely attached cells in AFM indentation experiments. Biophys J 2013;104:2123–31. 10.1016/j.bpj.2013.04.019.

[61] Weber A, Benitez R, Toca-Herrera JL. Measuring biological materials mechanics with atomic force microscopy – Determination of viscoelastic cell properties from stress relaxation experiments. Microsc Res Tech 2022;85:3284–95. 10.1002/jemt.24184.

[62] Abuhattum S, Mokbel D, Müller P, Soteriou D, Guck J, Aland S. An explicit model to extract viscoelastic properties of cells from AFM force-indentation curves. IScience 2022;25. 10.1016/j.isci.2022.104016.

[63] McCraw MR, Uluutku B, Solares SD. Linear Viscoelasticity: Review of Theory and Applications in Atomic Force Microscopy. Reports Mech Eng 2021;2:151–79. 10.31181/rme200102156m.

